# Biologically informed variational autoencoders allow predictive modeling of genetic and drug induced perturbations

**DOI:** 10.1101/2022.09.20.508703

**Authors:** Daria Doncevic, Carl Herrmann

## Abstract

Variational Autoencoders (VAE) have rapidly increased in popularity in biological applications and have already successfully been used on many omic datasets. Their latent space provides a low dimensional representation of input data, and VAEs have been applied for example for clustering of single-cell transcriptomic data. However, due to their non-linear nature, the patterns that VAEs learn in the latent space remain obscure. To shed light on the inner workings of VAE and enable direct interpretability of the model through its structure, we designed a novel VAE, OntoVAE (Ontology guided VAE) that can incorporate any ontology in its latent space and decoder part and, thus, provide pathway or phenotype activities for the ontology terms. In this work, we demonstrate that OntoVAE can be applied in the context of predictive modeling, and show its ability to predict the effects of genetic or drug induced perturbations using different ontologies and both, bulk and single-cell transcriptomic datasets. Finally, we provide a flexible framework which can be easily adapted to any ontology and dataset.

## Introduction

In recent years, deep learning (DL) has been widely used to analyze high-dimensional biological omics data, especially single-cell RNA-seq. In contrast to linear methods such as principal component analysis (PCA), non-linear models can capture more complex patterns in the data^1^. One prominent example of an unsupervised DL model that performs dimensionality reduction is the autoencoder (AE)^2^, which consists of two neural networks: an encoder, which compresses the data, and a decoder, which then aims at reconstructing the input data from this compressed representation that is also referred to as latent space. A more recent variant of the AE is the Variational Autoencoder (VAE)^3^ which learns a probability distribution over the latent vectors of the data and thus belongs to the class of generative models. Autoencoder based methods have successfully been applied in the context of cancer classification^4^, data integration^4^, data denoising^5^ and batch correction^6^, cell clustering^7^, multi-domain translation^8^ and prediction of effects of drug treatment on single-cells^9^. However, in contrast to PCA, these autoencoder based approaches lack interpretability as we cannot easily assign feature contributions to the latent vectors due to their intrinsically nonlinear nature.

Different approaches have already been used to tackle the problem of limited interpretability. Tybalt tries to extract a biologically meaningful latent space by examining how different latent vectors separate covariates and then investigating the associated gene weights of the latent vectors of interest in the one-layer decoder^10^. Other models modify the decoder to trade off reconstruction accuracy for interpretability. For example, in the scVI model, a linear decoder has been implemented to allow assignment of feature weights to the different latent vectors^1,11^. Another approach is the direct modification of the neural network structure through the incorporation of prior biological knowledge. In the VEGA and expiMap models, the authors used a one-layer, sparse decoder that connects the latent variables to a set of annotated genes, thus providing direct interpretability of the latent variables which can represent different biological entities such as pathways or transcription factors^12,13^. One limitation of these modelsis the simplicity of their structure, which does not allow the incorporation of more complex, hierarchical biological information.

Other approaches have been aiming at incorporating hierarchical biological networks into a neural network. Knowledge-primed neural networks (KPNNs), in which every node represents a protein or gene, and every edge a regulatory interaction, have been used to model T cell receptor stimulation^14^. DCell is structured according to subsets of Gene Ontology (GO) and has been used to predict growth rates in yeast and the impact of double-mutants on biological processes^15^. Gene Ontology Autoencoder (GOAE) implements GO terms in one hidden layer of encoder and decoder by partial connectivity to input and output layer, and has been used for clustering of single-cell RNA-seq data, however, as the ontology information is only used in one layer, GOAE can only model relationships between GO terms and genes, but not between parent and children GO terms^16^. Deep GONet is a neural network classifier that is imposing regularization on its weights to encourage the establishment of connections that mirror the GO directed acyclic graph (DAG), and has been used to combine cancer classification with biological explanations^17^. Its successor GraphGONet directly implements the GO structure without regularization^18^. However, to our knowledge, no attempts have yet been made to incorporate full hierarchical biological networks into a VAE in order to capture the different levels of description of biological processes in tasks that go beyond classification.

Here, we introduce OntoVAE (Ontology guided VAE), a novel flexible VAE architecture with a multi-layer, sparse decoder that allows for the incorporation of any kind of hierarchical biological information encoded as an ontology. OntoVAE provides direct interpretability in its latent space and decoder, as the activities of the neurons now correspond to activities of biological processes or phenotypes. Its efficient implementation allows users to consider thousands of terms and monitor their activity changes, without the need to pre-select specific processes. Importantly, OntoVAE can be used for predictive modeling. By modulating the values of input features *in silico* and then monitoring how these changes propagate through the network, OntoVAE can simulate the effects of drug treatment or genetic alterations. The investigation of subsequent alterations in the activation of hidden nodes representing processes or phenotypes allows to uncover complex genotype-phenotype relationships. Thus, we consider OntoVAE a powerful tool for *in silico* screening approaches.

## Results

### Architecture of OntoVAE

To make the latent space and decoder part of a VAE model biologically interpretable, we created a novel architecture that we named OntoVAE (Ontology guided VAE) where we implemented the latent space and decoder in a way that any biological ontology such as Gene Ontology (GO) or Human Phenotype Ontology (HPO) can be incorporated (Fig. 1a). Thus, every node in the latent space and every node in the decoder represents a term of the ontology, with root terms being located in the latent space and terms becoming more and more specific with progression through the decoder. Originally, every ontology has a common root node, but as a one-dimensional latent space would not be meaningful, we apply a trimming process to the ontology, where we remove the root and other very generic terms that have a high number of annotated genes. The resulting top terms after this trimming define the latent space. We furthermore remove very specific terms with too few annotated genes. Details on this pruning process can be found in *Methods* and Fig. S1.

**Fig 1.**
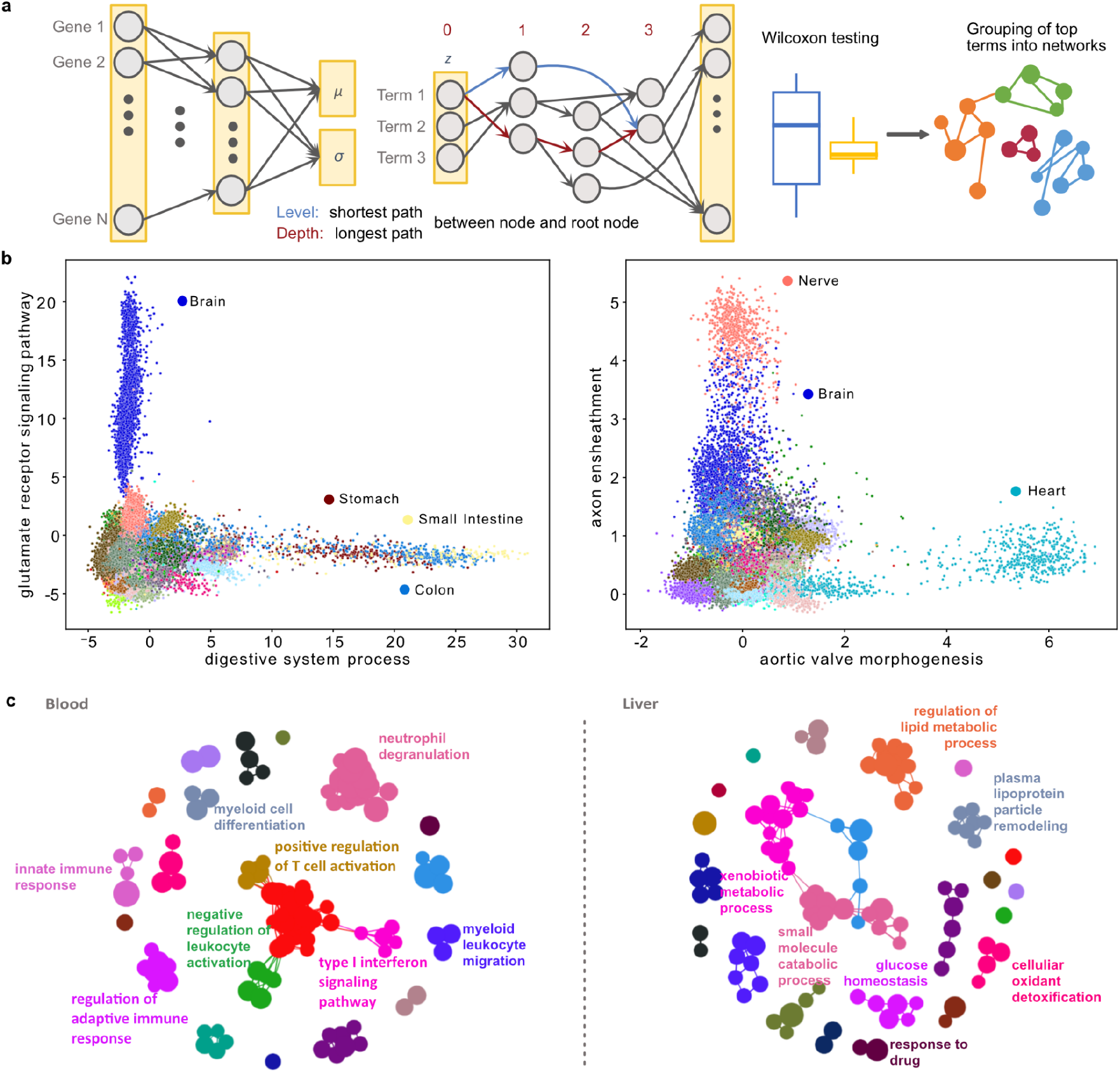
OntoVAE incorporates biological ontologies and provides pathway activities. **a** Schematic overview of OntoVAE. A non-linear encoder is coupled to a masked, multi-layer linear decoder. Both latent space and decoder are interpretable, as their structure and the modeled connections reflect a biological ontology. The latent space incorporates the root terms of the ontology, each layer in the decoder represents one depth level of the ontology, with ‘depth’ referring to the longest possible path between a node and a root node. Comparisons of node activation between samples is performed by Wilcoxon testing, and top terms for a group can be further summarized into networks based on their semantic similarities. **b, c** OntoVAE with GO-decoder has been trained on all tissue samples from GTEx. **b** Scatter plots showing example pathway activations for digestive system process and glutamate receptor signaling pathway (left), and aortic valve morphogenesis and axon ensheathment (right). **c** Example top term networks for Blood (left) and Liver (right).

The decoder is linear and sparse, meaning that no activation functions are used in the layers and connections are only modeled between parent and children ontology terms as well as between terms and annotated genes. Furthermore, we restrict all the weights in the decoder to be positive to preserve directionality in the model even when trained several times as has already been done in VEGA^12^. To model more complex relationships, each term of the ontology is modeled by three neurons.

### OntoVAE provides biological interpretability in latent space and decoder

We postulate that the activations of the neurons in the latent space and decoder can be directly interpreted as pathway or phenotype activities of the corresponding ontology. To demonstrate that, we incorporated the Gene Ontology (GO) into our model and trained it on the Genotype Tissue Expression (GTEx) bulk RNA-sequencing (RNA-seq) data^19–21^. We then retrieved the activities of all terms in the latent space and decoder for each sample and looked at some example terms to see if they were more active in the expected tissues. We found that, for example, *digestive system process* is especially active in stomach, colon and small intestine, *glutamate receptor signaling pathway* is especially active in brain, *axon ensheathment* is especially active in nerve and *aortic valve morphogenesis* is especially active in heart (Fig. 1b). In order for these results to be biologically meaningful, we should obtain similar results every time we train the same model with the same parameters. To confirm this, we trained the model twice using the same parameters, and using either one, two or three neurons per term, resulting in six trained models. For each GO term, we then calculated the Pearson correlation with itself when retrieving its pathway activities from two different models and found that the majority of correlations was higher than 0.95, thus ensuring the reproducibility of OntoVAE (Fig. S2a). To summarize our results in a tissue-centric manner, we performed Wilcoxon tests between all possible pairings of tissues at each node, and then assigned a rank to each term for each tissue (see Methods), allowing us to further group the terms that were most relevant for a given tissue into a network (Fig. 1a). As an example, we show the networks for blood and liver tissue (Fig. 1c). For blood, we find clusters related to *neutrophil degranulation, innate immune response, regulation of adaptive immune response,* and *myeloid cell differentiation.* For the liver, we find clusters related to *regulation of lipid metabolic process, response to drug, xenobiotic metabolic process,* and *glucose homeostasis.* This again confirms that our model generates meaningful biological results. The pathway activities for all GO terms as well as the networks for all tissues can be interactively explored in our web application (http://ontovaemodelexplorer.pythonanywhere.com/).

### OntoVAE can be used for in silico phenotype predictions of gene knockouts

Next, we investigated whether OntoVAE could be used for predictive modeling and simulate the outcome of a gene knockout. For this purpose, we performed an *in silico* knockout of the *Duchenne muscular dystrophy (DMD)* gene in the GTEx muscle samples. The *DMD* gene encodes for the protein dystrophin, which is located primarily in muscles and attaches the cytoskeleton to the extracellular matrix. Depletion of functional dystrophin protein due to a mutation in the *DMD* gene causes Duchenne Muscular Dystrophy (DMD), a genetic disease leading to muscle weakness and muscle degradation^22^.

The model was trained on all samples from all GTEx tissues. Then, the knockout was performed by setting the input value for the *DMD* gene to zero before applying the trained model on the muscle samples and obtaining the activity values^23^. We then used paired Wilcoxon tests at each node in latent space and decoder to compare which term activities significantly differed post-knockout (term-level analysis). We also computed paired Wilcoxon tests for each gene in the reconstruction layer to identify the most significantly affected genes, and performed overrepresentation analysis on the top 100 genes to further group them into GO terms (gene-level analysis) (Fig. 2a).

**Fig 2.**
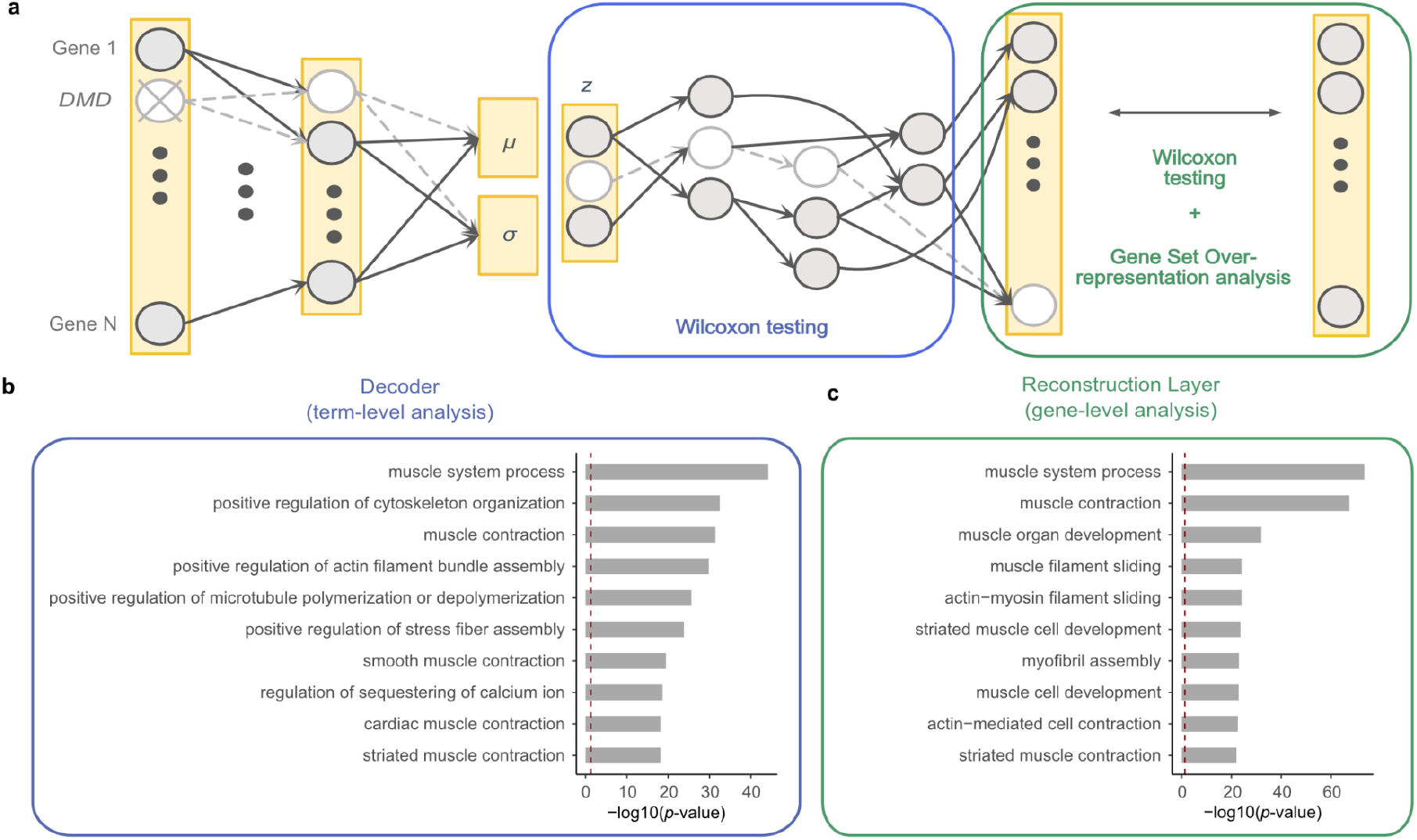
OntoVAE can predict phenotypic outcome of a gene knockout. OntoVAE has been trained with a GO-decoder on all GTEx tissue samples, knockouts have been performed in muscle samples only. **a** Schematic drawing of how the model can be used for *in silico* investigation of gene knockouts. Input value for a gene (here: *DMD)* is set to zero before running samples through the trained model and obtaining their activations at each node/term. Paired Wilcoxon tests are performed for all the terms in latent space and decoder (pre-knockout vs post-knockout) to identify the most affected terms (term-level analysis). Paired Wilcoxon tests can also be performed for all the genes in the reconstruction layer to identify the most affected genes, and these can then be further grouped into terms using gene set overrepresentation analysis (gene-level analysis). **b** Knockout of *DMD* in GTEx muscle samples. Barplot displays the ten most affected terms from the term-level analysis ranked by significance. **c** Knockout of *DMD* in GTEx muscle samples. Barplot displays the ten most affected terms from the gene-level analysis ranked by significance. **b,c** The red dotted line represents the significance threshold.

From the term-level analysis, we obtained terms that are highly related to *DMD* function, such as *muscle system process, positive regulation of cytoskeleton organization,* and *muscle contraction* (Fig. 2b). The gene-level analysis also yielded *DMD* related terms, such as *muscle system process, muscle contraction,* and *muscle organ development* (Fig. 2c). These results confirm our hypothesis that OntoVAE models meaningful relationships between the genes and can thus be used to predict the consequences of a gene knockout, with direct interpretability on a pathway level in latent space and decoder. We also wanted to show that these muscle-related terms are not the result of the fact that we used only GTEx muscle samples. Hence, as a negative control, we also performed the knockout for *SFSWAP,* a splicing factor, and *COX5A,* an enzyme of the mitochondrial respiratory chain, and obtained terms highly related to the respective gene (Fig. S3). All results of the paired Wilcoxon tests and the overrepresentation analysis can be found in Supplementary Table 2.

### OntoVAE can predict disease specific gene expression changes

Next, we asked whether OntoVAE could actually predict differential gene expression in the context of disease. As a use-case, we selected a form of muscular dystrophy, Limb-girdle muscular dystrophy, in order to evaluate to what extent our model can predict differentially expressed genes between disease and healthy control samples. We adapted our model to use the Human Phenotype Ontology (HPO)^24^, which consists of disease related terms, among them the term *Limb-girdle muscular dystrophy* (LGMD), a child node of the more generic term *Muscular dystrophy*. To predict the genes that most significantly influence the LGMD node in the decoder, we systematically performed a knockout for all genes one-by-one in the GTEx muscle samples by setting their input value to zero before passing the samples through the trained model, and then computed a paired Wilcoxon test at the LGMD node for each gene, comparing the activities at this node before and after knockout of the gene. For HPO, the sign of the influence of a gene on a term (either enhancing or inhibiting the phenotype) is not clearly defined. Therefore, we looked at both directionalities, and performed two separate one-tailed paired Wilcoxon tests for each gene, and then ranked the genes according to their *p*-values, ending up with two ranked lists: one list ranking the genes that downregulated the LGMD node upon knockout (predLGMD_dn) (Fig. 3a, Supplementary Table 3), the other list ranking the genes that upregulated the LGMD node upon knockout (predLGMD_up) (Fig. 3b, Supplementary Table 3). To verify the validity of these predictions, we performed a gene-set enrichment analysis (GSEA) using as a ground truth the differentially expressed genes in a recently published dataset of bulk RNA-seq carried out on muscle samples from LGMD patients (n=16) and healthy individuals (n=15)^25^, where we had determined the genes that were significantly up- (LGMD_up) or downregulated (LGMD_dn) in patients compared to age-matched controls (Supplementary Table 3). For both lists (predLGMD_up and predLGMD_dn), we found a significant enrichment of LGMD_dn (Fig. 3a,b), confirming that OntoVAE is able to predict differentially expressed genes between disease and control. We fused the genes from the leading edges of the two GSEA analyses, and checked the overlap of those predicted genes with LGMD_dn, and with genes directly annotated to the term LGMD in the HPO (Fig. 3c). We found an overlap of 147 and 10, respectively. Next, we were interested to verify if the 10 genes directly annotated to LGMD ended up in the leading edge of the prediction as a result of their direct annotation to LGMD. Hence, for these 10 genes, we trained 10 new models, in which we removed the link to LGMD for one of the 10 genes. We then performed the knockout again to see if the genes would still end up in their respective leading edge. Of those 10 genes, 7 had significantly downregulated the activity at the LGMD node, and were still located in the leading edge of predLGMD_dn after their link to LGMD had been removed (Fig. 3d). The remaining 3 genes had significantly upregulated the activity at the LGMD node, and one of them was still in the leading edge of predLGMD_up after link removal (Fig. 3e). This analysis confirms that OntoVAE can tolerate missing prior information to a certain degree, as for the majority of the genes it recovered their importance in absence of direct annotation. It also confirms that OntoVAE learns complex relationships that go beyond the direct annotation of a gene to a specific term.

**Fig 3.**
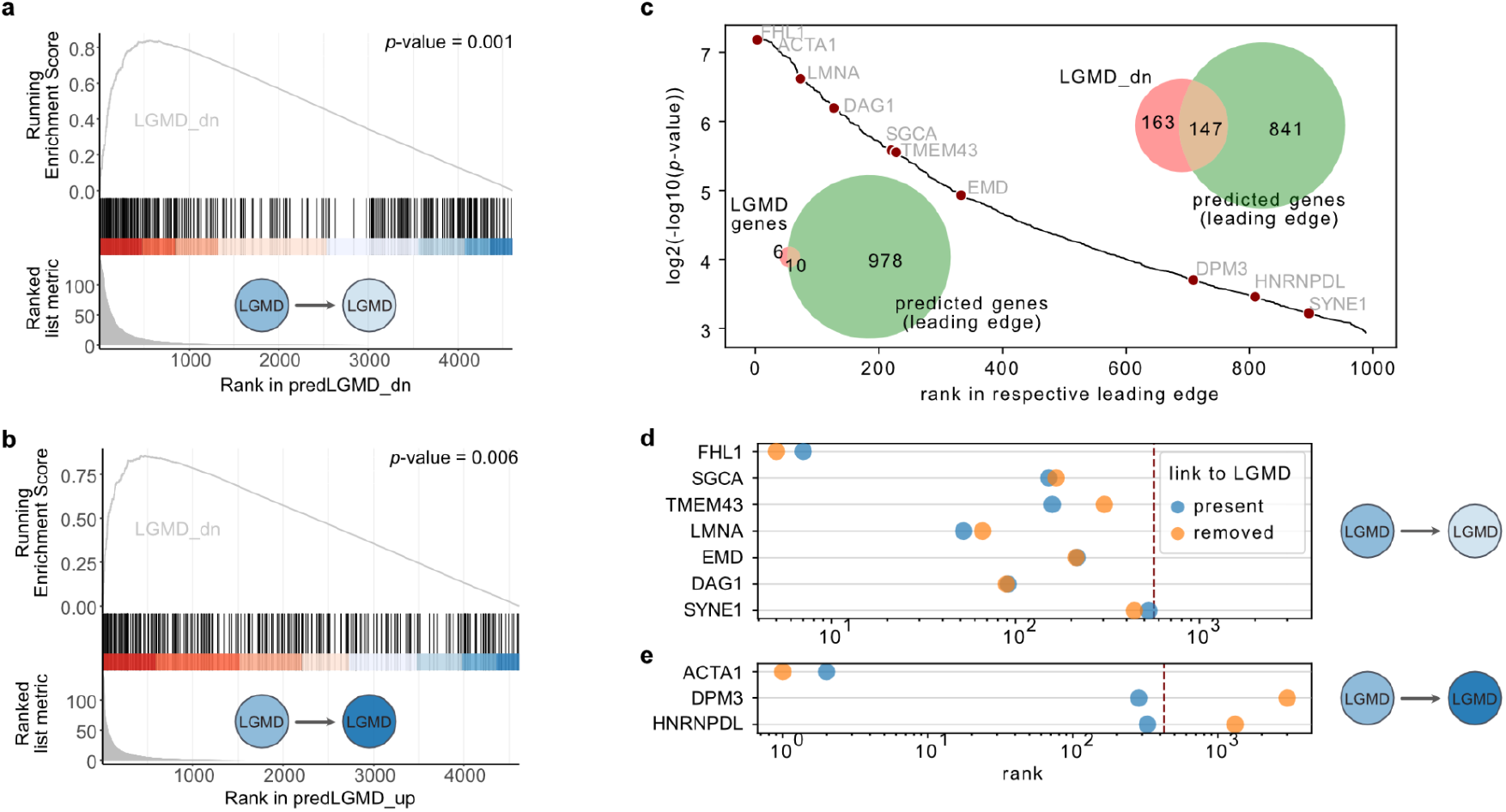
OntoVAE can predict differential gene expression in disease. OntoVAE has been trained with HPO-decoder on all tissue samples from GTEx, in-silico knockouts for all genes were performed in muscle samples only. All genes were then ranked according to their impact on the node corresponding to Limb-girdle Muscular Dystrophy (LGMD), and GSEA was performed with the genes that were up-/down-regulated in patients versus control (LGMD_up/LGMD_down) in an external validation LGMD dataset. **a** GSEA results using ranking of genes that significantly downregulated the LGMD node in the decoder (predLGMD_dn). **b** GSEA results using ranking of genes that significantly upregulated the LGMD node in the decoder (predLGMD_up). **c** Leading edges of both GSEAs were fused (predicted genes). Venn diagram in the top right corner shows overlap between the predicted genes and LGMD_dn. Venn diagram in the bottom left corner shows overlap between predicted genes and genes directly annotated to the term LGMD in HPO. Hockey stick plot displays the ranking of the predicted genes, the 10 genes directly annotated to LGMD in HPO are labeled. **d, e** For each of the 10 genes, a new model was trained where the direct link of the gene to the LGMD term had been removed. Plots show the ranking of the 10 genes before (blue circle) and after (orange circle) removing the link to LGMD. **d** Genes that were found in the leading edge of **a**, red dotted line indicates leading edge cutoff. **e** Genes that were found in the leading edge of **b**, red dotted line indicates leading edge cutoff.

### OntoVAE can predict treatment effects in silico

Finally, we wanted to investigate the predictive power of OntoVAE in the context of drug treatment, so we shifted back to GO to focus on interferon response. For this purpose, we used the peripheral blood mononuclear cell (PBMC) dataset from Kang *et al.,* in which PBMCs from lupus patients were treated with interferon (IFN)-ß (a type I IFN) and single-cell RNA-seq was performed^26^. A UMAP representation of this dataset reveals clustering by treatment condition and cell type (Fig. 4a).

**Fig 4.**
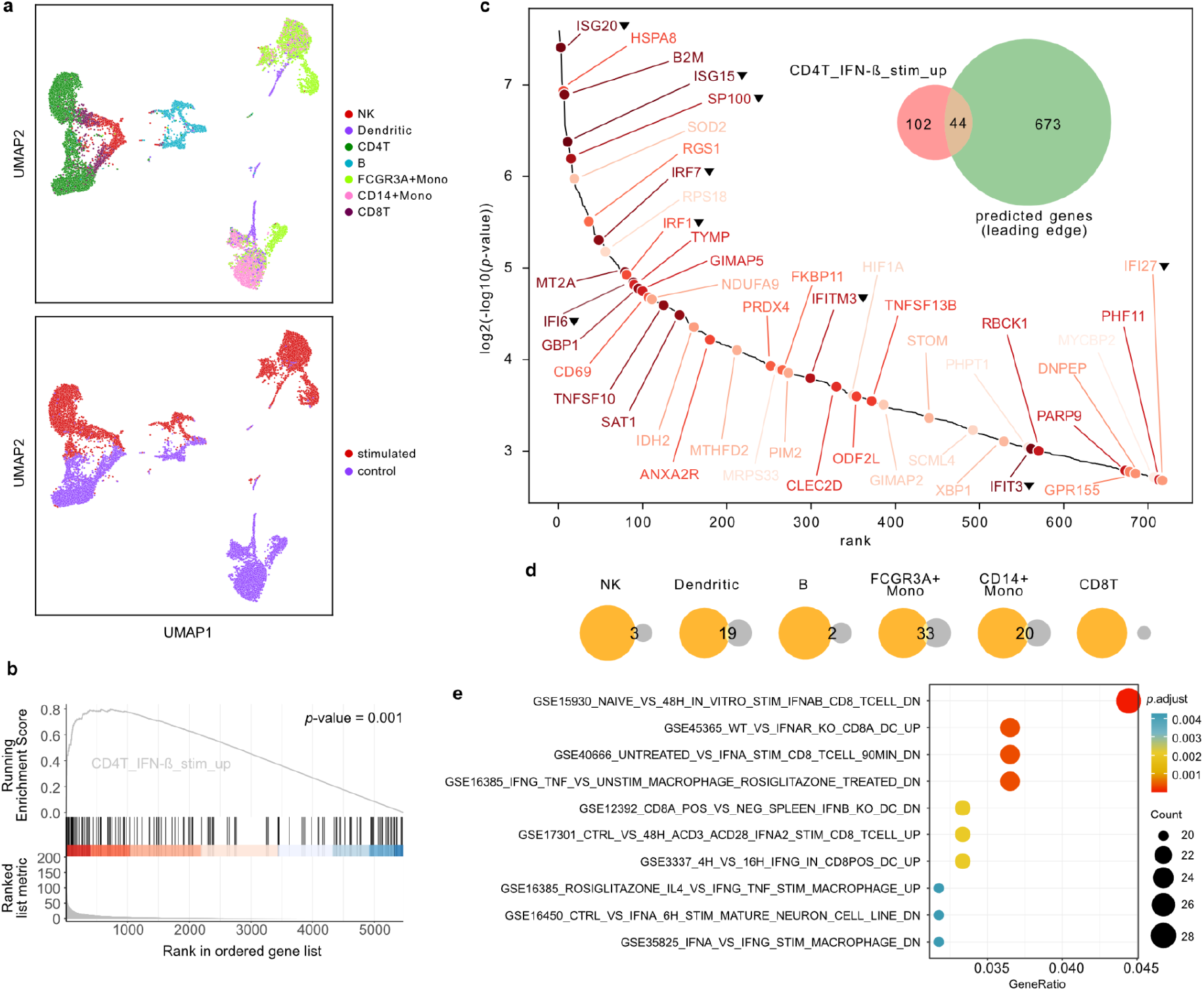
OntoVAE can predict interferon treatment response. **a** UMAP plot shows the PBMC dataset colored by cell type (top panel) and treatment (bottom panel). **b, c** Prediction of interferon treatment response by *in silico* stimulation of genes. All in-silico stimulated genes were ranked according to their impact on the activity of the *type I interferon signaling pathway* node. **(b)** GSEA was performed with the genes that were upregulated in interferon-ß (IFN-ß) treated CD4T cells versus control CD4T cells in the original dataset (CD4T_IFN-ß_stim_up). Based on GSEA, genes from the leading edge (termed ‘predicted genes’) were selected. **(c)** Venn diagram displaying the overlap between the predicted genes (green circle) and CD4T_IFN-ß_stim_up (red circle). The hockey stick plot is displaying the predicted genes sorted by significance, the 44 genes overlapping with CD4T_IFN-ß_stim_up are labeled. Darker label color indicates higher significance of this gene in CD4T_IFN-ß_stim_up, a black triangle next to the label indicates that the gene is directly annotated to the term *type I interferon signaling pathway* or to one of its descendant terms. **d, e.** Further investigation of predicted genes that do not overlap with CD4T_IFN-ß_stim_up (green circle in Venn diagram **c** without intersection). Venn diagrams are showing the overlap of these genes (orange circles) with the genes that are upregulated upon IFN-ß treatment in the other cell types of the PBMC dataset (gray circles) (**d**). Dotplot is showing results from a gene set overrepresentation analysis using interferon related terms from MSigDBs C7 immunologic signature gene sets (**e**).

For our analysis, we looked at CD4T cells, as they represent the largest population from the dataset. First, we performed differential gene expression analysis between stimulated and unstimulated CD4T cells and identified 146 genes that are significantly upregulated in the IFN stimulated CD4T cells (CD4T_IFN-ß_stim_up, Supplementary Table 4). We used these as ground truth or ‘reference genes’ in our analysis. We then trained our model with a GO-decoder on all unstimulated cells, and then performed one-by-one *in silico* stimulation of all input genes by setting their expression value to a higher value (see Methods). To investigate which genes had the highest influence on the node in the GO-decoder corresponding to *type I interferon signaling pathway,* we computed a paired Wilcoxon test between the activations at this node before and after stimulation for each gene in the CD4T cells (Supplementary Table 4). The genes were then ranked according to their paired Wilcoxon *p*-value, and gene set enrichment analysis (GSEA) was performed with the gene set CD4T_IFN-ß_stim_up, yielding a significant *p*-value of 0.001 (Fig. 4b) and therefore confirming that OntoVAE can predict interferon response related genes. We had a closer look at the genes from the leading edge of the GSEA, which we refer to as ‘predicted genes’, and 44 of them overlapped with our ground truth CD4T_IFN-ß_stim_up (Fig. 4c), among them some of the most significant reference genes such as *ISG15, IFI6, ISG20,* and *B2M.* Notably, of those 44 genes, 35 genes are not directly annotated to the term *type I interferon signaling pathway* or any of its descendant terms (Fig. 4c), showing that the model can also recover genes that are not directly linked to the node of interest. Also, among those genes, some had very low expression or were expressed in very few of the unstimulated cells in the PBMC dataset, which comprised our training data, but could still be recovered by OntoVAE (Fig. S4). We then further analyzed the 673 predicted genes that were not overlapping with the reference genes, and found substantial overlap with the genes that were differentially upregulated upon IFN-ß treatment in some of the other cell types of the PBMC dataset, especially FCGR3A+ monocytes, CD14+ monocytes, and dendritic cells (Fig. 4d). We also performed gene set overrepresentation analysis (ORA) with the interferon related gene sets from MSigDBs C7 immunologic signature gene sets (Fig. 4e), finding significant enrichment of gene sets that were upregulated in CD8T cells or a mature neuron cell line upon IFN treatment, or downregulated in dendritic cells after IFN receptor knockout (first, second, third and ninth set in Fig. 4e). Taken together, these results confirm that OntoVAE generates meaningful results through *in silico* prediction of treatment response.

## Discussion

In this work, we designed OntoVAE, a novel VAE, in whose latent space and decoder any hierarchical biological network can be incorporated that has the structure of a directed acyclic graph (DAG) and can be trimmed to have a reasonable number of root nodes that represent the latent space variables. We applied the model on different ontologies, Gene Ontology (GO) and Human Phenotype Ontology (HPO), and on a bulk RNA-seq dataset (GTEx) as well as on a single-cell RNA-seq dataset (PBMC). Using GTEx and GO, we demonstrated that the model generates reproducible results and outputs meaningful pathway activities in latent space and decoder. Furthermore, we applied OntoVAE in two scenarios of predictive modeling: genetic alterations and drug treatments. Again using GTEx and GO, we showed that we obtain meaningful terms in the decoder and reconstruction layer when simulating gene knockouts. Using GTEx and HPO, we showed how to systematically perform an *in silico* knockout of all genes to find which genes had the strongest influence on the disease LGMD, and validated our findings in an external RNA-seq dataset of LGMD. Using PBMC and GO, we showed how the model, trained on unstimulated data only, is still able to predict interferon response by systematically performing *in silico* stimulation of all genes. One interesting and central finding is the fact that the model is capable of predicting influential genes on processes or phenotypes that go beyond the ones directly annotated to the term in question, indicating that the model is capable of learning more complex gene-term relationships in a data driven way. Previous models have tackled the question of interpretable drug-response prediction, for example the CPA model based on autoencoders coupled to discriminators, which provides interpretability in terms of covariate influence, such as species or treatment dose^27,28^. Other models, such as Dcell^15^ and a recently published model to predict gene expression from transcription factor activities^23^ have also employed synthetic knockouts. The recently published expiMap model, based on a VAE with an interpretable latent space, can also predict drug response and learn new relationships based on prior knowledge such as pathway structure^13^. However, unlike other models that limit the number of biological terms under scrutiny or use a single-layer linear decoder, our model is capable of encoding thousands of terms without the need for preselection and maintaining the hierarchical information contained in the ontology. In addition, our model extends to synthetic over-expression of genes to simulate drug-induced activation. As such, it uses a similar approach as the light-up procedure implemented in DeepVAE, but at the level of the input gene expression instead of the nodes of the hidden layer^29^.

One limitation of our model is that we only model one type of relationship between terms, namely the ‘is a’ relationship, as there is no straight-forward way to model different kinds of connections. By enabling more complex and diverse relationships between nodes, our current model could incorporate knowledge-graphs as a decoder in a similar fashion as we did for ontologies as an alternative to graph-neural networks that are commonly used^30^. Additionally, as we use a hard-coded decoder, the model has no possibility of correcting the prior. However, we also demonstrate that in some cases, the model can recover missing prior information, for example missing gene annotations to terms, suggesting that it is capable of learning complex relationships. Nevertheless, the quality of the results will heavily depend on the quality of the used prior information and adding flexibility in the connections of the decoder along the lines proposed by Rybakov et al.^31^ or as implemented in DeepGONet^17^ might be an interesting option to cope with incomplete annotation. For biological ontologies, one has to keep in mind that some branches are more explored than others, some terms are more central than others, with more incoming and outgoing connections, and also the genes are annotated to different numbers of terms. To which extent these parameters influence the results remains to be investigated. Furthermore, for some ontologies such as HPO where many annotations are based on SNPs, it is challenging to interpret the results as annotated genes can have a positive or negative effect on the term, whereas our model only allows for positive weights in the decoder in order to preserve the direction when interpreting pathway activities. With our systematic knockout approach, we were only able to predict genes that were downregulated in LGMD patients, thus, further investigation is needed here. We also want to point out that the model cannot distinguish between causal and indirect relationships.

In summary, OntoVAE can be adapted to any ontology and dataset and used to compute pathway activities and to predict disease or treatment induced changes in gene expression. In this manuscript we focused on single perturbations. In future work, it will be interesting to also investigate the effect of multiple perturbations at a time, and see if synergistic effects can be recovered by OntoVAE. We envision that our method will be a useful first step in identifying sets of target genes or pathways which can then be further validated experimentally.

## Methods

### Variational Autoencoder (VAE)^3^

The VAE is a probabilistic deep learning model consisting of two coupled, but independently parameterized neural networks: an encoder which learns a probability distribution over a compressed representation of the high-dimensional input data, which we refer to as latent space, and a decoder which tries to reconstruct the original input from this latent space representation after sampling from the learnt distribution. More formally, the encoder part, or recognition model, yields a posterior distribution *qθ{z/x),* with *θ* being the learnable parameters of a neural network, *x* the input data and the latent space. The decoder part, or generative model, yields a likelihood distribution *p_ϕ_(^x/z^*), with *Φ* being the learnable parameters of a neural network^32^. During the training process, both parts are jointly optimized using stochastic gradient descent. As computation of the posterior is intractable, the optimization objective of the VAE, like of other variational inference methods, is the *evidence lower bound* (ELBO)^12,32^:

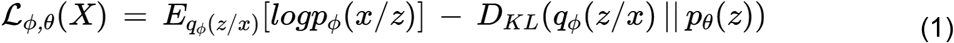

The first part of equation (1) is trying to optimize the likelihood of the observed data under a given model. The second part of equation (1) is the Kullback-Leibler (KL) divergence, which determines the divergence of the approximate posterior from the true posterior. In practice, the true posterior *pθ(z)* is often modeled as a multivariate normal distribution, thus, the KL divergence has a closed-form solution^3^. The reparameterization trick is applied during model training to enable standard backpropagation^3^.

### OntoVAE architecture

OntoVAE is a modified VAE whose latent space and decoder are implemented so that they can incorporate any directed acyclic graph (DAG) such as a biological ontology (Fig. 1a). For this manuscript, we used Gene Ontology (GO) and Human Pathway Ontology (HPO). A DAG is suitable for incorporation into a deep learning model due to its hierarchical structure and some of its main properties, such as the absence of connected loops and the possibility of a child term to have multiple parent terms (unlike a tree). Note the definition of ‘level’ and ‘depth’ here, with ‘level’ referring to the shortest possible path between a node and a root node, and ‘depth’ referring to the longest possible path between a node and a root node. For ontology incorporation into the model, we define the latent space as the root layer (depth 0), and each layer of the decoder corresponds to one depth level of the ontology, meaning that terms of the ontology that have the same depth will be located in the same layer. Thus, the most generic terms are located in the latent space, and terms become more and more specific with progression through the decoder until reaching the reconstruction layer, which is made up of genes. Note that, although we only used gene expression data in this project, in principle, any type of data can be used as input and output that can be mapped to the terms of the ontology used, such as DNA methylation or SNP data. A DAG usually has one root node, but because a one-dimensional latent space would not be meaningful, we apply a trimming process to our ontologies before incorporation, where the most generic terms, with too many annotated genes, as well as the most specific terms, with too few annotated genes, are removed (Fig. S1). Thresholds for pruning can be chosen by the user.

The decoder of OntoVAE is linear, multi-layered and sparse, as it only models connections between parent and child terms of the ontology as well as between terms and annotated genes. In contrast to a standard VAE where connections are only present between neighboring layers, OntoVAE also has to model connections between non-neighboring layers as well as between any layer and the reconstruction layer, as terms in each depth level have genes annotated to them. This is achieved by a successive layer concatenation process when passing samples through the model, where at each step, the values from the previous decoder layer are concatenated to the current one. Furthermore, at each step, a binary mask is used to set the weights of non-existent connections to zero.

We postulate that the activation value we retrieve from a neuron located in the latent space or decoder of the model when running samples through the trained model corresponds to the pathway activity of the term represented by this neuron. To preserve directionality in the pathway activities, we restrict the weights of our decoder to be positive. Furthermore, we use three neurons per term to compensate for the orientation of the ontology in the model, with information flowing from parent to children terms, and reduce correlations between siblings that share the same parent nodes and would therefore receive the same input if only one neuron per term was used (Fig. S2). Thus, in order to obtain the pathway activity for one term, we take the average of the activities of the three neurons representing the term.

### Ontologies and datasets

The preprocessed ontologies and datasets used in the manuscript are provided under https://figshare.com/projects/OntoVAE_Ontology_guided_VAE_manuscript/146727.

Ontologies were downloaded in obo format and parsed with the function *get_godag()* from the Python package *goatools* (v 1.0.15).

#### Gene Ontology (GO)^19^

Ontology was downloaded from *geneontology.org* (go-basic.obo, release date: 2021-02-01) together with the GAF annotation file (goa-human.gaf, version 2.1). We filtered everything for the ‘Biological Process’ sub-ontology and kept only ‘is a’-relationships between terms. We furthermore trimmed the ontology using a bottom threshold of 30 and a top threshold of 1000 (Fig. S1), ending up with 3083 GO terms and 19,469 annotated genes when using HGNC symbols, and 3245 GO terms and 19,387 annotated genes when using Ensembl IDs. HGNC symbol to GO term annotations were extracted from the GAF file. To obtain Ensembl ID to GO term annotations, Uniprot IDs from the GAF file were converted to Ensembl using a mapping file from *ftp.uniprot.org* (idmapping_selected.tab, download: April 2021).

#### Human Phenotype Ontology (HPO)^24^

The ontology was downloaded from *hpo.jax.org* (hp.obo, release date: 2022-04-14) together with an annotation file (genes_to_phenotype.txt) containing the mapping of HPO IDs to genes. We applied the same preprocessing and trimming steps as for GO, but this time using thresholds of 10 and 1000, and ended up with 4525 HPO terms and 4774 annotated genes represented by HGNC symbols.

#### Genotype Tissue Expression (GTEx) dataset^21^

GTEx bulk RNA-seq data from human tissues was retrieved through the R package *recount3* (v 1.1.8), which is communicating with recount3, a database of uniformly reprocessed RNA-seq datasets^33^.

#### Limb-girdle muscular dystrophy (LGMD) dataset^25^

Bulk RNA-seq data sequenced from muscle samples of LGMD patients and healthy individuals were downloaded from Gene Expression Omnibus (accession number: GSE202745).

#### Peripheral blood mononuclear cells (PBMC) dataset^26^

The preprocessed single-cell RNA-seq dataset of interferon (IFN)-ß treated PBMCs was downloaded through the github repository of VEGA (https://github.com/LucasESBS/vega-reproducibility).

### Model implementation and training

The OntoVAE framework is written in pytorch. All OntoVAE models in this manuscript have been trained with a batch size of 128, a dropout of 0.2 in the hidden layer of the encoder, a dropout of 0.5 on the latent space layer, and a weighting coefficient of 1e-4 on the Kullback Leibler loss. Models were usually trained over 300 epochs, and the model with the best validation loss was used. AdamW was used as the optimizer together with a learning rate of 1e-4. The dimensions of latent space and decoder are always defined by the used ontology. Three neurons were used per ontology term. The following pretrained models are provided under https://figshare.com/projects/OntoVAE_Ontology_guided_VAE_manuscript/146727: 1. GO with Ensembl IDs trained on GTEx bulk RNA-seq data downloaded through recount3, 2. HPO with Gene Symbols trained on GTEx bulk RNA-seq data downloaded through recount3, 3. GO with Gene Symbols trained on unstimulated cells from the PBMC dataset.

### Retrieval and comparison of pathway activities

We retrieve the activities at each node when running samples through a pre-trained model by attaching pytorch hooks to each layer. To compare pathway activities between groups of samples, we performed two-tailed Wilcoxon tests using the *stats.ranksums* function from the *scipy* package (v 1.5.2) with a *p*-value cutoff of 0.05 at each node in the decoder.

### Creation of Gene Ontology Networks

For the creation of tissue-specific GO networks for the GTEx dataset (Fig. 1c), we performed Wilcoxon tests at each node/for each term for all possible two pairs of tissues, and called it a hit if a term was significantly more active in one tissue compared to another (*p*-value < 0.05). We then counted the number of hits per tissue and used this as a measure to rank all tissues for one term, with the tissue with most hits receiving rank 1 for this term, the tissue with second most hits receiving rank 2 and so on. For a given tissue, the terms were then sorted again based on their rank, the number of hits, and the median of the Wilcoxon test statistic. We then took the top 100 terms of the tissue and clustered them based on their Wang semantic similarities^34^. Thus, each node in the network corresponds to one GO term, the size of a node reflects the number of genes associated with a term. If all terms in a cluster had a common ancestor, this ancestor term was chosen as cluster representative. Otherwise, the term with most annotated genes was chosen as cluster representative. GO networks for all tissues can be explored in our Dash based web application (http://ontovaemodelexplorer.pythonanywhere.com/. Supplementary table 1 contains the rankings of the GO terms for all tissues.

### Simulation of genetic and drug induced perturbations

All simulations in this study were carried out in a single-gene context, meaning that we only perturbed the expression of one gene at a time. For the study of genetic perturbations, *in silico* gene knockouts have been performed by setting the input value for a gene to zero before running data through the trained model and retrieving pathway activities (Fig. 2a). For the study of drug induced perturbations, *in silico* stimulation of genes was performed by increasing their input value prior to running data through the trained model.

#### GTEx muscle + GO

Knockout of *DMD, COX5A,* and *SFSWAP* was performed in the 881 muscle samples.

#### GTEx muscle + HPO

Systematic knockout of all 4774 genes annotated to HPO was performed in the 881 muscle samples.

#### CD4T cells (unstimulated) from PBMC dataset + GO

Systematic stimulation of all 5472 genes annotated to GO and expressed in the unstimulated cells of the PBMC dataset was performed by setting their input value to 8, as 6.8 was the maximum expression value found in the unstimulated cells.

### Perturbation analysis (paired Wilcoxon test)

For analysis of perturbation consequences, one-tailed paired Wilcoxon tests were performed between the same set of samples pre- and post perturbation using the function *stats.wilcoxon* from the *scipy* package (v 1.5.2).

#### GTEx muscle + GO

Tests were performed at each node in latent space and decoder to identify terms significantly downregulated upon knockout (term-level analysis, Fig. 2b). Tests were also performed for each gene in the reconstruction layer to identify genes significantly downregulated upon knockout (gene-level analysis, Fig. 2c).

#### GTEx muscle + HPO

Tests were performed specifically at the decoder node *Limb-girdle muscular dystrophy* in both directions, to identify genes that significantly up- or downregulated the activity of this node. Genes were then ranked according to their *p*-value.

#### CD4T cells (unstimulated) from PBMC dataset + GO

Tests were performed specifically at the decoder node *Type I interferon signaling pathway* to identify genes that significantly upregulated the activity of this node. Cells that originally had zero expression of a particular gene were not included in the test. The number of cells that was included per gene can be found in Supplementary Table 4. Genes were then ranked according to their *p*-value.

### Differential gene expression analysis (DGEA)

#### PBMC dataset

DGEA was performed between IFN stimulated and unstimulated cells for each cell type separately. Only genes were included that could also be mapped to GO. We performed two-tailed Wilcoxon tests using the *stats.ranksums* function from the *scipy* python package (v 1.5.2) and corrected for multiple testing with the function *stats.multitest.fdrcorrection* from the python package *statsmodels* (v 0.12.0). For each cell type, we then selected genes significantly upregulated upon IFN treatment, using an FDR threshold of 0.1. Gene set upregulated in CD4T cells was termed CD4T_IFN-ß_stim_up.

#### LGMD dataset

DGEA was performed between muscle samples of patients (n=42) and controls (n=33) using the R package *DESeq2* (v 1.26.0). Eight patient samples and one control sample had been excluded as outliers prior to analysis after PCA inspection of the dataset. We used a significance threshold of 0.1 for the adjusted *p*-value and kept only genes that could be mapped to HPO. Genes upregulated in LGMD muscles compared to control were termed LGMD_up, genes downregulated in LGMD muscles compared to control were termed LGMD_dn.

### Gene set enrichment analysis (GSEA)

GSEA was performed with the R package *clusterProfiler* (v 3.14.3).

### Gene set overrepresentation analysis (ORA)

ORA was performed with the R package *clusterProfiler* (v 3.14.3).

## Data availability

All datasets used for analysis in this manuscript are publicly available and listed in the Methods section. Further data generated in this manuscript such as pretrained models, preprocessed ontologies or curated gene lists can either be found on figshare https://figshare.com/projects/OntoVAE_Ontology_guided_VAE_manuscript/146727 or in the Supplementary material or can be made available by the authors upon request.

## Code availability

The package for preprocessing ontologies and training OntoVAE models is available at https://github.com/hdsu-bioquant/onto-vae. Code to generate the results of the manuscript is provided under https://github.com/hdsu-bioquant/OntoVAE_manuscript. A web application to interactively explore all results related to Fig. 1 of the manuscript can be accessed under: http://ontovaemodelexplorer.pythonanywhere.com/.

## Acknowledgments

DD and CH are supported by the e:Med project COMMITMENT (Grant 01ZX1904D) from the German ministry for research and education (BMBF). This project is further supported by the DFG grant *Identification of the molecular origins of comorbidity in COVID-19 patients* (DFG HE 7201/3-1) to CH. We thank Andrés Quintero and Emanuel Schwarz for fruitful discussions and helpful comments on the manuscript and Steeve Boulant for discussions on the interferon application. CH acknowledges financial support by IPMB, University Heidelberg.

## Author contributions

DD and CH conceived the idea. DD implemented the model and carried out the analyses. CH supervised the project. DD and CH wrote the manuscript.

---

**Supplementary Table 1** Results of GO term ranking for the different tissues of GTEx.

**Supplementary Table 2** Results of Wilcoxon testing to analyze *DMD, SFSWAP* and *COX5A* knockout.

**Supplementary Table 3** LGMD_dn and LGMD_up from LGMD bulk RNA-seq dataset and results of paired Wilcoxon testing to identify genes that up- or downregulate LGMD activity upon knockout.

**Supplementary Table 4** CD4T_IFN-ß_stim_up from PBMC single-cell RNA-seq dataset and results of paired Wilcoxon testing to identify genes that upregulate IFN activity upon stimulation.

**Fig S1.**
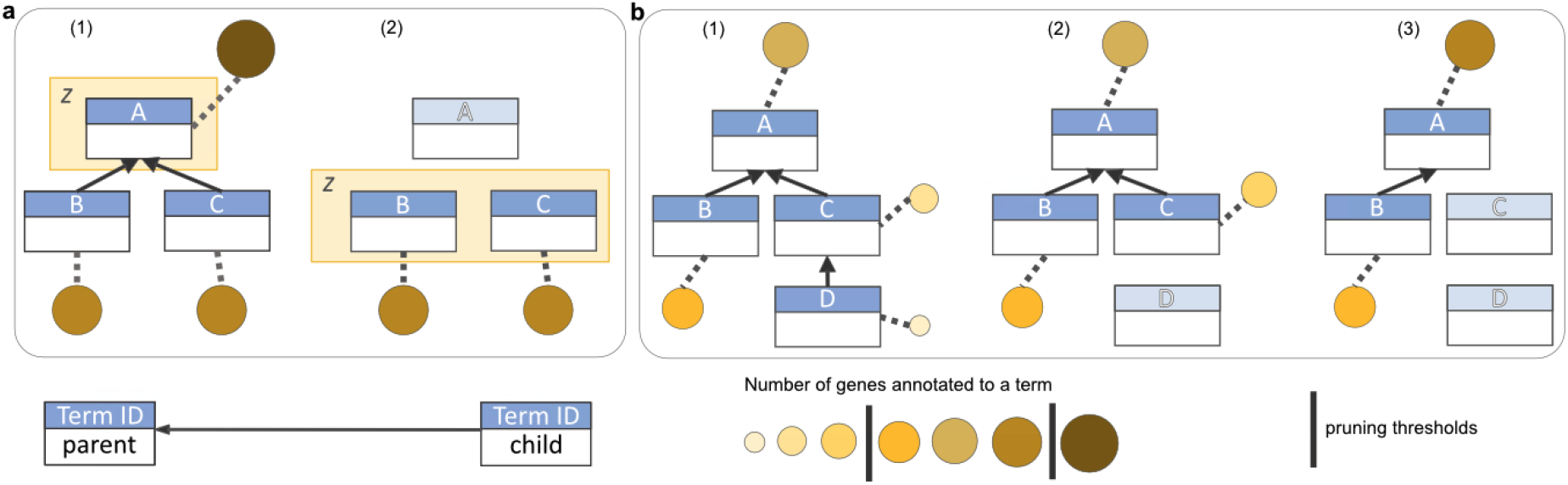
Threshold-based ontology pruning for integration into OntoVAE. **a** Top pruning: term A is a root term located in the latent space *z* and has more genes annotated to it than the top pruning threshold (1). Thus, term A will be removed and its children terms B and C will become part of the latent space (2). **b** Bottom pruning: term D has less genes annotated to it than the bottom pruning threshold (1), so its genes are transferred to its parent term C, and D is removed (2). Term C still has less genes annotated to it than the bottom threshold, so its genes are transferred to its parent term A, and term C is pruned (3). Pruning stops here, as the number of genes annotated to A lies within the thresholds.

**Fig S2.**
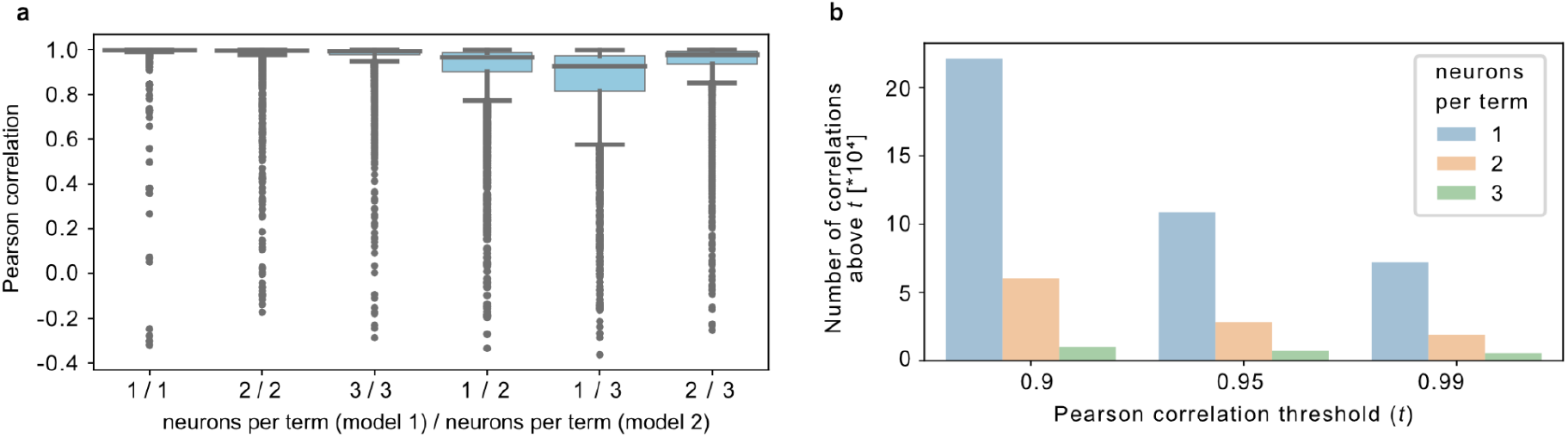
Fig. S2 OntoVAE generates reproducible results. Different models were trained with GTEx data and GO with the same parameters, but using 1, 2, or 3 neurons per term, respectively. **a** Boxplots showing the Pearson correlations for the same GO terms between different models. **b** Barplots showing the number of pairwise Pearson correlations above a certain threshold for 1, 2, and 3 neurons per term.

**Fig S3.**
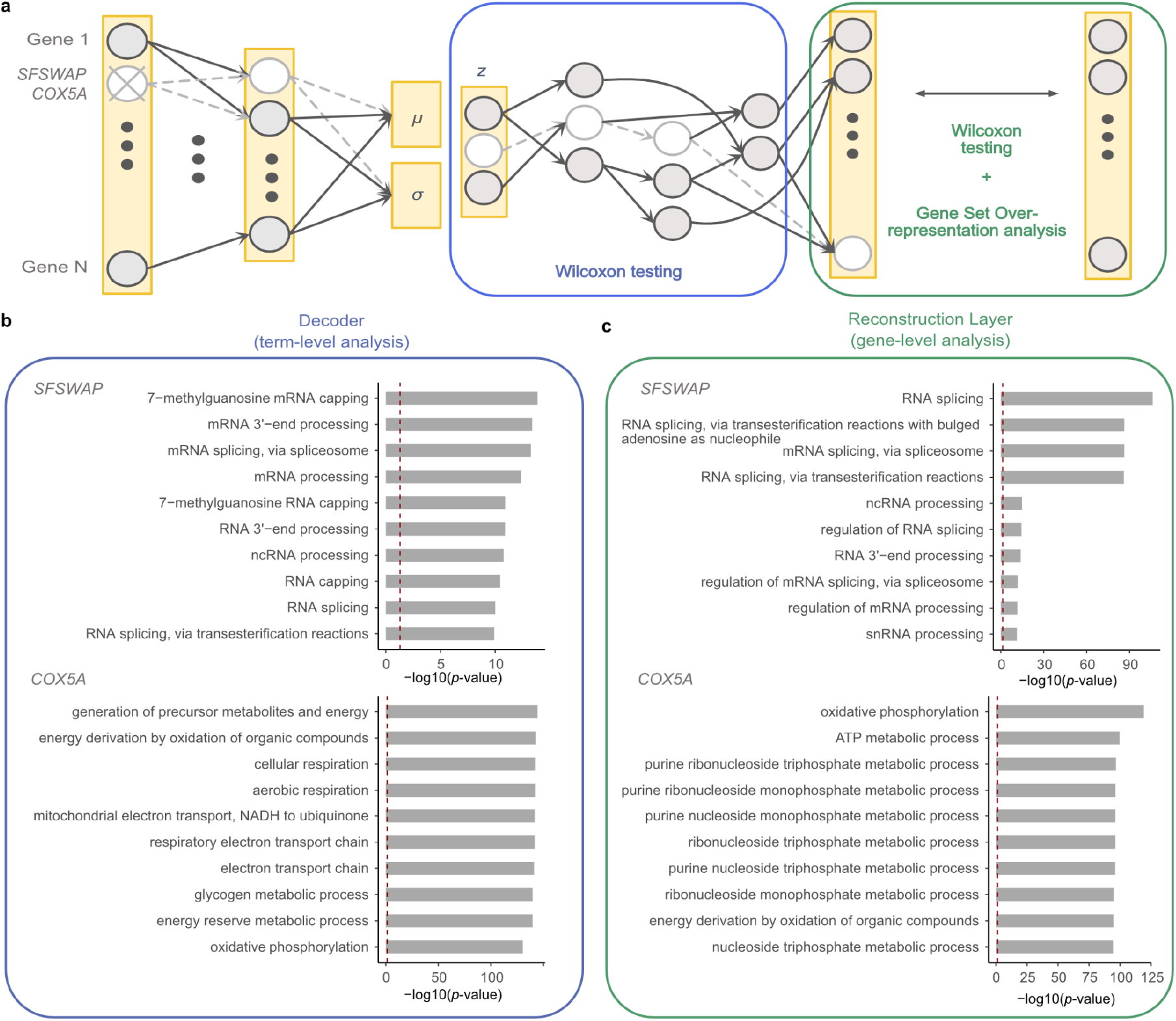
OntoVAE can predict phenotypic outcome of a gene knockout (here: *SFSWAP* and *COX5A*. **a, b, c** as in Fig. 3. **b** Barplots displaying the term-level analysis results for *SFSWAP* and *COX5A.* **c** Barplots displaying the gene-level analysis results for *SFSWAP* and *COX5A.*

**Fig S4.**
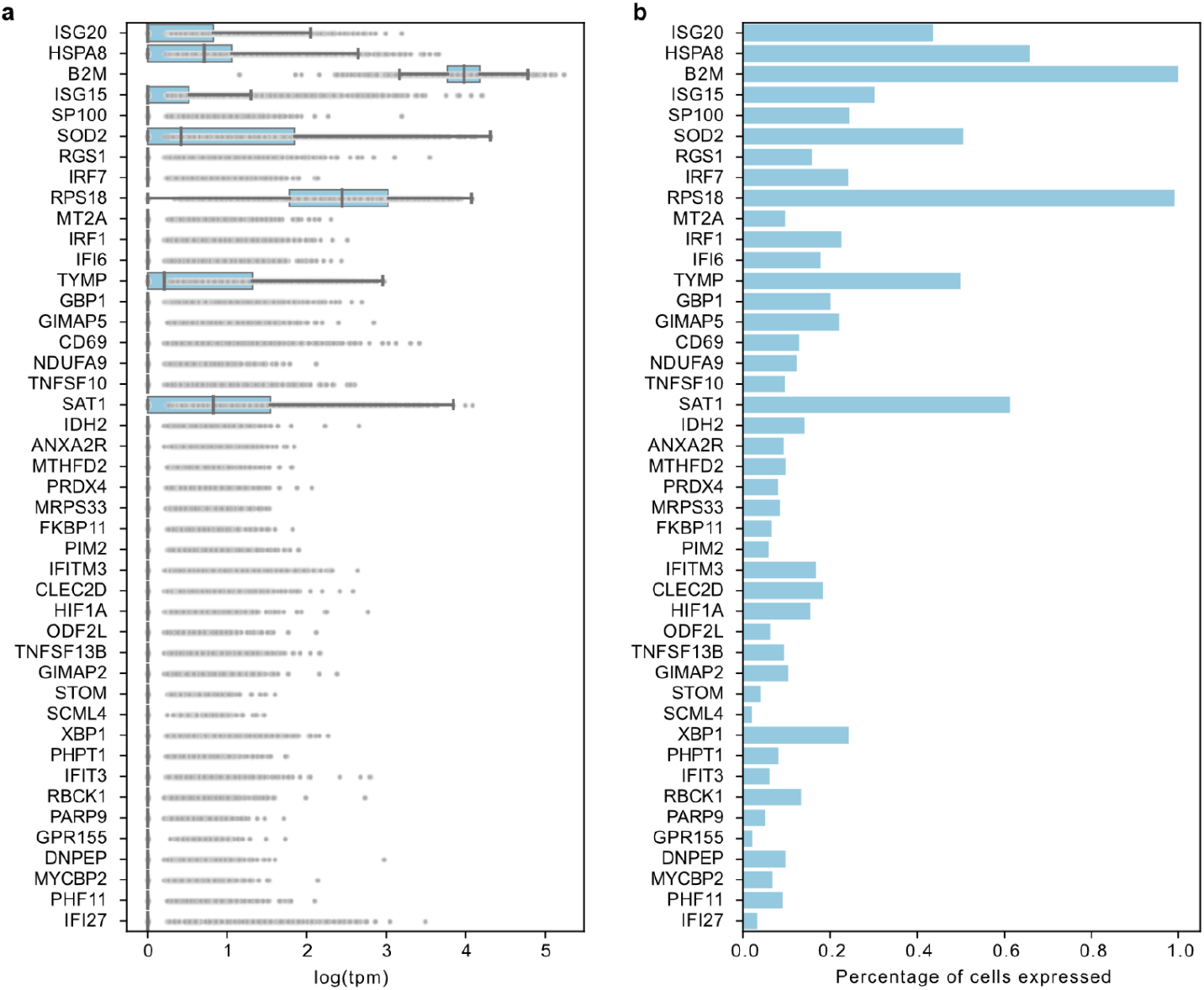
OntoVAE can predict genes with low expression in training data. Displayed genes are found in the intersection of genes upregulated upon IFN-ß treatment in CD4T cells and genes predicted by OntoVAE to influence the node *type I interferon signaling pathway.* **a** Boxplots show the expression of the 44 genes in the training data, **b** barplots show what percentage of cells in the training data expresses the gene. Note that the training data consists of all unstimulated cells from the PBMC dataset.

